# Estimating Error Models for Whole Genome Sequencing Using Mixtures of Dirichlet-Multinomial Distributions

**DOI:** 10.1101/031724

**Authors:** Steven H. Wu, Rachel S. Schwartz, David J. Winter, Donald F. Conrad, Reed A. Cartwright

## Abstract

**Motivation:** Accurate identification of genotypes is critical in identifying de novo mutations, linking mutations with disease, and determining mutation rates. Because de novo mutations are rare, even low levels of genotyping error can cause a large fraction of false positive de novo mutations. Biological and technical processes that adversely affect genotyping include copy-number-variation, paralogous sequences, library preparation, sequencing error, and reference-mapping biases, among others.

**Results:** We modeled the read depth for all data as a mixture of Dirichlet-multinomial distributions, resulting in significant improvements over previously used models. In most cases the best model was comprised of two distributions. The major-component distribution is similar to a binomial distribution with low error and low reference bias. The minor-component distribution is overdispersed with higher error and reference bias. We also found that sites fitting the minor component are enriched for copy number variants and low complexity region. We expect that this approach to modeling the distribution of NGS data, will lead to improved genotyping. For example, this approach provides an expected distribution of reads that can be incorporated into a model to estimate de novo mutations using reads across a pedigree.

**Availability:** Methods and data files are available at https://github.com/CartwrightLab/WuEtAl2016/.

## 1 Introduction

Identifying genotypes from next-generation sequencing (NGS) data is an important component of modern genomic analysis. Accurate genotyping is key to identifying sequence polymorphisms, detecting de novo mutations, linking genetic variants with disease, and determining mutation rates (Awadalla *et al.*, 2010; Goldstein *et al.*, 2013; Koboldt *et al.*, 2013; Peng *et al.*, 2013; Sayed *et al.*, 2009). Inaccurate genotyping affects the accuracy of identifying de novo mutations, as true mutations are rare compared to errors in sequencing and downstream analyses. Because putative SNPs are typically validated using another sequencing technology, high false positive rates increase the effort required for validation.

However, estimating genotypes from NGS data can be computationally and statistically challenging. A typical NGS experiment generates millions of short read fragments, 100–650 base-pairs in length, that are aligned to a reference genome. If the only error was due to sampling, NGS methods would produced data that was perfectly representative of the underlying genotype; the base calls for homozygous sites would be identical, and the base calls for heterozygous sites would follow a 1:1 binomial distribution. For this reason, genotyping and variant calling software initially used binomial distributions to model heterozygous base-counts (Li *et al.*, 2008a,b; Goya *et al.*, 2010; Cartwright *et al.*, 2012).

However, there are at least three experimental processes that are thought to affect the ratio of the alleles. (1) During library preparation, variation in amplification rates might cause some chromosomes to be replicated more than others (Heinrich *et al.*, 2012). This variation is especially a concern if there is little starting material. (2) NGS technologies can introduce sequencing errors into sequencing reads. Error rates are on the order of 0.1–1% per base-call. While this may seem small, 0.1% error is equivalent to sequencing the wrong human genome, and 1% is equivalent to sequencing a chimpanzee instead of a human (Fox *et al.*, 2014; Wall *et al.*, 2014). (3) Bioinformatic methods that assemble reads with respect to a reference can misplace reads and penalize non-reference alleles (Degner *et al.*, 2009; Krawitz *et al.*, 2010). Together these processes can shift the mean and increase the variance of sequencing read count distributions. These processes do not affect all parts of the genome equally. The genomic context of a site, including the presence of nearby indels, structural variants, or low complexity regions, influences the probability that reads generated from a given site will be subject to these processes (Malhis and Jones, 2010; Li, 2014). Thus, it is possible for both homozygotes and heterozygotes to have an intermediate ratio of two alleles, and it can be difficult to accurately identify genotypes using a binomial distribution (Malhis and Jones, 2010).

Current approaches to calling genotypes from NGS data deal with the issues described above to some degree. The increased variance and skewed allele ratios expected to be produced from mismapped reads can be partially controlled by including mapping quality data in a genotype-calling procedure. In the simplest approaches, reads with low quality scores are removed from an analysis. In Bayesian approaches to genotype calling, read quality data may included when calculating genotype probabilities (e.g. Li *et al.*, 2009b).

The increased variance expected to be caused by library preparation, sequencing, and errors in mapping reads to a reference genome can be accommodated by modeling read-counts as coming from a beta-binomial distribution (Ramu *et al.*, 2013). The beta-binomial distribution acts as an over-dispersed binomial, allowing the excess variance to be handled in a standard statistical framework. The Dirichlet multinomial distribution (DM) is the general case of the beta-binomial, allowing for overdispersion and modeling of more than two outcomes. The DM has been used to model allele counts and the frequency of multi-allelic genotypes within tumor samples (Josephidou *et al.*, 2015; Tvedebrink, 2010; Muralidharan *et al.*, 2012). Such genotype calling procedures can be combined with machine learning algorithms that attempt to differentiate between true variants and those caused by sequencing artifacts (DePristo *et al.*, 2011).

In this study, we evaluate the hypothesis that finite mixtures of Dirichlet-multinomial can produce a better fit to the underlying error processes. The mismatch between observed and expected read distributions created by the processes described above contributes to observed false positive single nucleotide polymorphism (SNP) discovery rates of 3 to 15% (Harismendy *et al.*, 2009; Farrer *et al.*, 2013). We posit that improving the expected read distribution (i.e. the model) will reduce false positives. As a first step toward this goal in this study we model the distribution of base counts produced from NGS using a mixture of Dirichlet-multinomial distributions (MDMs). Furthermore, we model sites that are the most likely to be true heterozygotes (rather than false positives). By modeling these sites exclusively we determine the true effect of variation in amplification rates, error, and mismapping.

Fitting MDMs to sequencing data improves existing genotyping methods in two important ways. First, we can account for the context-dependent nature of genotyping errors by allowing multiple DM distributions, each with different parameter values, to be estimated for a given dataset. Additionally, by using more than two categories we explicitly model the presence of bases that are neither reference nor the likely alternative allele at a given site. Thus, we are able to directly estimate the probability of sequencing errors in a given DM model.

We first demonstrate the value of our approach by fitting MDMs to sequencing data derived from a haploid human cell line (CHM1). The MDM produces a superior fit to this data compared to other models, showing that even relatively simple genetic datasets can be the result of heterogeneous processes, and thus benefit from a mixed-model approach. We then fit MDMs to diploid data generated by the 1000 Genomes Project (1000 Genomes Project Consortium *et al.*, 2010, 2015). For these datasets, the MDM also improves the fit compared to other models. Most interestingly, when we limit our data to “true” heterozygotes, most of the data fit a binomial distribution, rather than the expectation of over-dispersed, reference-biased data. Using a model that fits the data should lead to significant improvements in genotyping, which in turn should lead to fewer candidate mutations that are found to be inaccurate when validated in a lab.

## 2 Methods

### 2.1 Data

We extracted datasets from two types of data. The first dataset is the haploid human sequence from a hydatidiform mole cell line (CHM1hTERT SRR1283824 from SRP017546). We refer to this dataset as the CHM1 dataset in this paper.

Second, we obtained Illumina sequences from the 1000 Genomes Project for three individuals, a woman (NA12878) and both of her parents (NA12891 and NA12892). Sequencing was repeated for these individuals in different years using different technologies (2010, 2011, 2012, 2013). The 2010 dataset was generated during pilot 2 studies with short and variable read length between 36bp and 76bp. The 2011 and 2012 dataset were not part of a release. They contain the same sequencing data, 101bp reads, with slightly different bioinformatics. The 2013 dataset was part of the phase 3 release, and was sequenced without PCR, with longest read length of 250bp (1000 Genomes Project Consortium *et al.*, 2010, 2015).

We refer to this dataset as the CEU dataset. If the release year is appended, for example CEU2013, then we refer to the specific release in 2013. For each of these five datasets (CHM1 and each of the four releases of CEU), we analyzed two genomic regions, the whole chromosome 21 and a subregion of chromosome 10, from positions 85534747 to 135534747, which is approximately the same size as chromosome 21 (48 million base pairs).

For CHM1, we called genotypes by first obtaining allele counts for each base at each site using the mpileup function in SAMTools v1.2 (Li *et al.*, 2009a; Li, 2011) and the human reference genome (Genome Reference Consortium human genome build 37). We then used BCFTools v1.2 (Li *et al.*, 2009a; Li, 2011) to identify sites called as homozygous reference. For each of these sites we calculated the frequency of the reference allele and the frequency of all non-reference alleles (error). We filtered this dataset based on the read depth for each site: we removed sites with read counts of less than 10 or greater than 150. The upper limit is calculated using a method based on Warr *et al.* (2015), which takes the median plus twice the standard deviation of coverage per site; sites with zero coverage are ignored. Sites with high numbers of reads are likely in copy number variable genes that have aligned to a single region of the genome. In sites with extreme coverage, apparent heterozygotes are more likely to be due to paralogs rather than variation within a gene. The low read filter limits the data to calls with enough coverage to provide a reasonably accurate call and proportion of reads for each base. We refer this dataset as the full dataset (FD).

We also focused on modeling sites that were very likely to be homozygous reference sites in order to understand what the distribution of reads for those sites looks like. To generate our reference dataset (RD), we removed sites for which less than 80% of the reads contained the reference allele. The cutoff point was picked after examining the empirical distributions; there were only a few sites for which 50% to 80% of the reads contained the reference allele.

For the CEU data, we obtained allele counts as above for all three individuals. We then called genotypes as above on NA12878 (the daughter of the trio) and by using the BCFtools trio caller with the data from all three individuals. We limited the dataset we used for subsequent analyses to sites that were found by both methods. Sites that were found only in trio caller, but not in the individual caller were likely identified by the pedigree with limited data for the daughter; thus, these low coverage sites were not included in subsequent analysis. We removed sites with read counts of less than 10 or greater than 150, as for CHM1. We call this the Potential Heterozygote (PH) dataset. For each of these sites we calculated the frequency of the reference allele, the frequency of the alternate allele, and the frequency of any other alleles (error). We compared the frequencies of each allele category (reference, alternate, error) for each possible genotype combination. Because we found no differences in frequencies for different genotypes, all subsequent analyses were only performed on the general reference-alternate-error dataset.

We created an additional dataset by removing sites from the PH dataset not found to be heterozygous by the 1000 Genomes Project (1000 Genomes Project Consortium *et al.*, 2010, 2015). We then discarded sites for which the alternate allele differed from the one previously identified by the 1000 Genomes Project. We call this the True Heterozygote dataset (TH).

Because the CHM1 dataset was larger than the CEU dataset, we randomly subsampled the CMH1 dataset to have an approximately equal number of sites (40,000 sites) as the CEU dataset.

### 2.2 Model fitting and parameter estimation

We fit seven models to each CHM1 dataset, and eight to each CEU dataset. The models included a multinomial, a multinomial with reference bias (CEU only), Dirichlet multinomial (DM) and mixtures of DM (MDM) distributions with various number of components, ranging from two to six. We estimated the parameters and calculated the genotype likelihood for each model.

The genotype likelihood measures the likelihood of a sample’s genotype, *G*, given a set of base-calls, *R*, and is proportional to the probability of observing *R* if the genotype was *G*, i.e. *L*(*G*|*R*) ∝ *P*(*R*|*G*). We derived genotype likelihoods using MDM distributions. The DM is a compound distribution generated when a Dirichlet distribution is used as a prior for the probabilities of success of a multinomial distribution: *p* ∼ Dirichlet(α) and *x* ∼ Multinomial(N,p) where α is a vector of concentration parameters,p is a vector of proportions, *x* is a vector of counts, and *N* is the sample size. After integrating out *p*, the resulting probability mass function can be trivially expressed as a product of ratios of gamma functions:

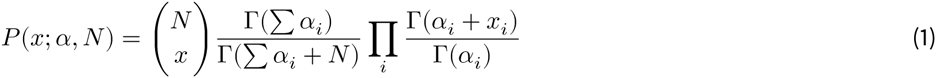

where *i* is one of the nucleotide, ∑*x_i_* = *N* and *α_i_* > 0. Furthermore,

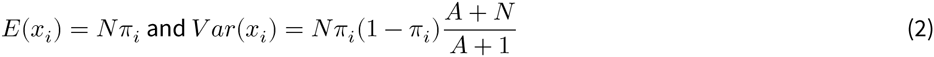

where *A* = ∑*α_i_* and 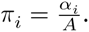.

It is helpful to reparameterize the distribution by letting 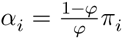, where 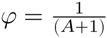 represents the pairwise correlation between samples. As a result, 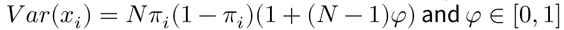 is a parameter controlling the amount of excess variation in the Dirichlet multinomial. When φ = 0, the DM reduces to a multinomial. Thus the Dirichlet multinomial can be interpreted as an over-dispersed multinomial distribution. As φ increases the distribution becomes more overdispersed, and at φ = 1 every observation contains only one type of nucleotide.

For a single-component Dirichlet-multinomial, we estimated the maximum-likelihood model starting with a method-of-moments estimation and optimizing using the Newton-Raphson method. For MDM, the maximum-likelihood estimated was computed using an EM algorithm and random starting points. This procedure was repeated 1000 times to search for the global maximum likelihood estimation.

For the Dirichlet-multinomial distribution, we estimated φ as a measure of the overdispersion of the data. In addition, we estimated *ρ*, the mixing proportion of sites belongs to each Dirichlet-multinomial component. For the homozygous CHM1 dataset, we estimated the proportion of the reference allele and error alleles for each model or model component. For each CEU dataset we estimated the proportion of the reference allele, alternate allele, and the error alleles. The reference and alternate allele were determined by SAMTools.

To determine the optimal number of components in the MDM model we calculated the Akaike information criterion (AIC) and Bayesian information criterion (BIC) for each dataset. The model with the lowest AIC or BIC is considered the best fitting model; when the best-fit model differs for AIC and BIC we use the BIC-selected model. The AIC and BIC for each model are calculated by the following formulas:

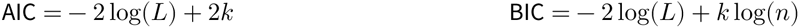

where *L* is the maximum likelihood estimate from the model, *k* is the number of free parameters, and *n* is the number of sites in the dataset.

### 2.3 Visualizing model fit

We visualized the fit of the data to each model and compared the fit between models using quantile-quantile (QQ) plots. The QQ plot plots the quantile of the observed read count frequencies against the quantile of the expected read count frequencies. Parameters estimated from the EM were used to estimate the expected read count frequencies by Monte Carlo simulation with 100 replicates. Two QQ plots, one for the reference allele and one for the error term, were used to illustrate the fit of models for the CHM1 dataset. Three separate QQ plots, one each for reference, alternate, and error terms, were used for the CEU datasets.

### 2.4 Frequency of sites in copy number variable regions and low complexity regions

In order to explore the composition of each component, the likelihood of every site was calculated under each component of the model in each of the CEU datasets. The likelihood for each component was evaluated using the parameters estimated from the EM. By comparing the likelihood between all components, each site was assigned to the component with the highest likelihood.

We extracted all the known CNV sites for NA12878 in the CEU dataset (Mills *et al.*, 2011). We calculated the number and proportion of sites belong to the known CNVs for the major and minor components (combined) for the best fit model for each of the eight datasets. We used a Fisher’s exact test to determine whether there is a significant difference between the proportion of CNVs in each component. We extracted all known LCRs in the human reference from the UCSC Genome Bioinformatics with the Table Browser data retrieval tool (Karolchik *et al.*, 2004), and repeated the same analyses performed for the CNV regions. We calculated the number and proportion of LCRs in the major and minor components for the best fit model and used a Fisher’s exact test to determine whether there is a significant difference between these two components.

## 3 Results

### 3.1 Haploid dataset — Homozygous reference

We examined two genomic regions from the CHM1 (haploid human cell line) dataset: all of chromosome 21 and part of chromosome 10. For each region we further split sites into two subsets. The full dataset (FD) contains base calls that were only filtered to exclude regions with unusually high coverage and contains both haploid reference and haploid non-reference sites. In order to exclude haploid non-reference sites, we remove sites containing less than 80% reference base calls, generating the reference dataset (RD).

#### Best fitting models

Using the expectation maximization (EM) algorithm, we fit seven models to each genomic region in each dataset: a multinomial, a DM, and MDM models with two to six components. The addition of model components increased the likelihood of the model for all cases (supplementary tables). We used Bayesian information criterion (BIC) to penalize overfitting and select the best-fitting model; the best model for each dataset was the two component MDM. Similar results were obtained when using Akaike information criterion (AIC) (see supplement). Each site was assigned to one component based on its maximum likelihood calculated on the parameters estimated by EM. In all cases one component contained a substantial majority of sites (approximately 75% of sites for the reference dataset from chromosome 21 and 95% of the sites for other datasets). We will refer to the component to which the highest proportion of sites was assigned as the “major component” and all other components as “minor components”.

In the DM distribution, the overdispersion parameter, φ, describes the degree to which the expected variance of a given distribution is greater than that of a corresponding multinomial. This parameter can take values between 0 and 1, with 0 being identical to the multinomial and 1 being completely overdispersed. For the FD, the major component had relatively little overdispersion (φ = 0.00252 and 0.00415 for chromosome 21 and chromosome 10 respectively). For this dataset, the minor component displayed strong overdispersion (φ = 0.892 and 0.948). When we fitted MDMs to the reference datasets (in which sites with a high proportion of non-reference alleles were removed) the major component had less overdisperson compared to the full dataset (φ = 0.00 and 0.00269). The minor components of models fitted to this dataset were slightly overdispersed (φ = 0.0153 and 0.0475) (Table 1 and supplementary tables).

**Table 1.**
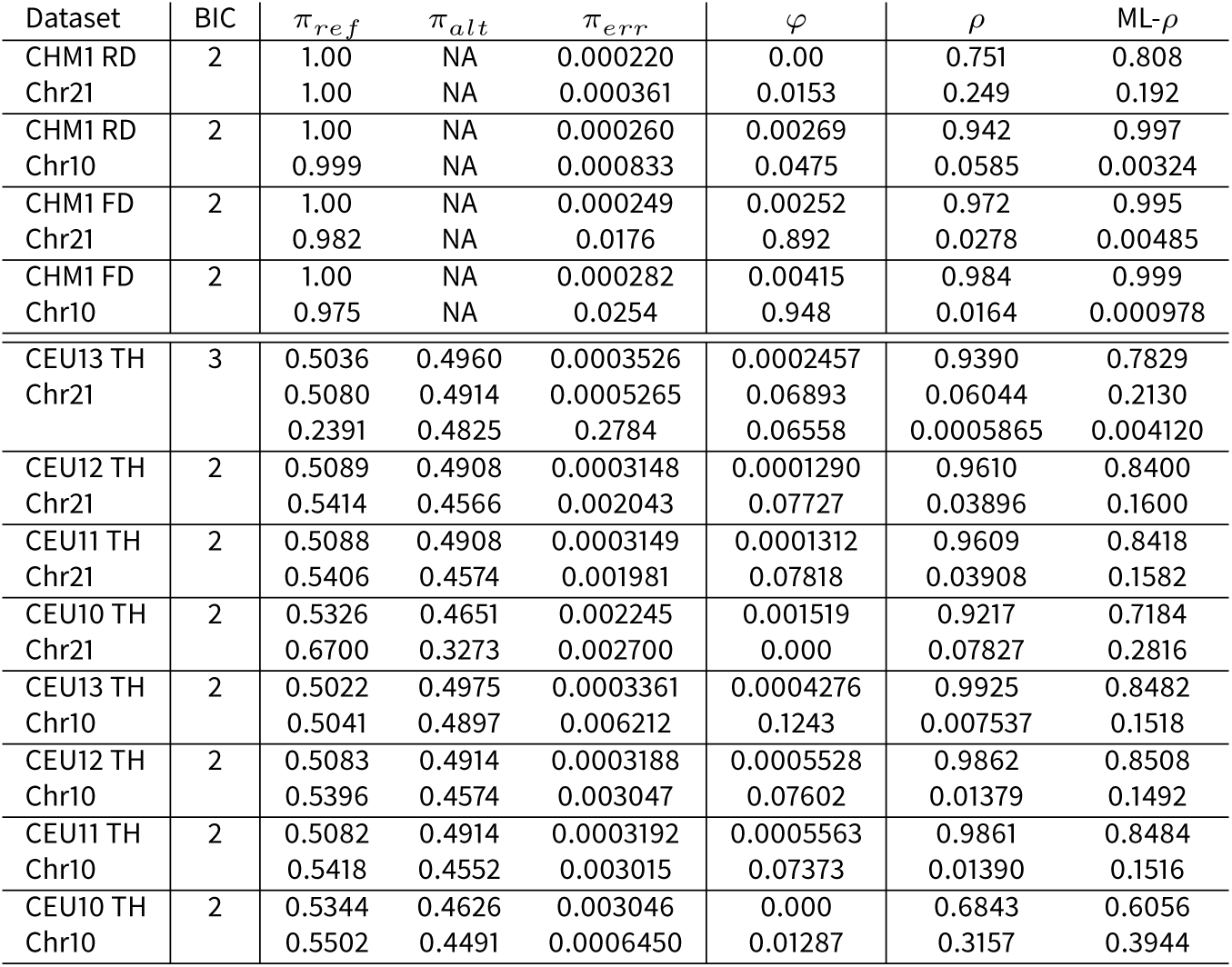
The number of components in the best MDM model according to BIC values for CHM1 and CEU dataset and parameters estimated for the best BIC model. π_ref_, π_alt_, and π_err_ is the proportion of the reference, alternative and error term respectively. *φ* is the overdispersion parameter. When *φ* = 0, the DM reduces to a multinomial. As φ approaches 1, the distribution is completely overdispersed. *ρ* is the percent of sites in each component. ML-*ρ* is the percent of sites assigned to each component using the likelihood.

| Dataset | BIC | $\pi_{ref}$ | $\pi_{alt}$ | $\pi_{err}$ | $\varphi$ | $\rho$ | ML-ρ |
| --- | --- | --- | --- | --- | --- | --- | --- |
| CHM1 RD | 2 | 1.00 | NA | 0.000220 | 0.00 | 0.751 | 0.808 |
| Chr21 |  | 1.00 | NA | 0.000361 | 0.0153 | 0.249 | 0.192 |
| CHM1 RD | 2 | 1.00 | NA | 0.000260 | 0.00269 | 0.942 | 0.997 |
| Chr10 |  | 0.999 | NA | 0.000833 | 0.0475 | 0.0585 | 0.00324 |
| CHM1 FD | 2 | 1.00 | NA | 0.000249 | 0.00252 | 0.972 | 0.995 |
| Chr21 |  | 0.982 | NA | 0.0176 | 0.892 | 0.0278 | 0.00485 |
| CHM1 FD | 2 | 1.00 | NA | 0.000282 | 0.00415 | 0.984 | 0.999 |
| Chr10 |  | 0.975 | NA | 0.0254 | 0.948 | 0.0164 | 0.000978 |
| CEU13 TH | 3 | 0.5036 | 0.4960 | 0.0003526 | 0.0002457 | 0.9390 | 0.7829 |
| Chr21 |  | 0.5080 | 0.4914 | 0.0005265 | 0.06893 | 0.06044 | 0.2130 |
|  |  | 0.2391 | 0.4825 | 0.2784 | 0.06558 | 0.0005865 | 0.004120 |
| CEU12 TH | 2 | 0.5089 | 0.4908 | 0.0003148 | 0.0001290 | 0.9610 | 0.8400 |
| Chr21 |  | 0.5414 | 0.4566 | 0.002043 | 0.07727 | 0.03896 | 0.1600 |
| CEU11 TH | 2 | 0.5088 | 0.4908 | 0.0003149 | 0.0001312 | 0.9609 | 0.8418 |
| Chr21 |  | 0.5406 | 0.4574 | 0.001981 | 0.07818 | 0.03908 | 0.1582 |
| CEU10 TH | 2 | 0.5326 | 0.4651 | 0.002245 | 0.001519 | 0.9217 | 0.7184 |
| Chr21 |  | 0.6700 | 0.3273 | 0.002700 | 0.000 | 0.07827 | 0.2816 |
| CEU13 TH | 2 | 0.5022 | 0.4975 | 0.0003361 | 0.0004276 | 0.9925 | 0.8482 |
| Chr10 |  | 0.5041 | 0.4897 | 0.006212 | 0.1243 | 0.007537 | 0.1518 |
| CEU12 TH | 2 | 0.5083 | 0.4914 | 0.0003188 | 0.0005528 | 0.9862 | 0.8508 |
| Chr10 |  | 0.5396 | 0.4574 | 0.003047 | 0.07602 | 0.01379 | 0.1492 |
| CEU11 TH | 2 | 0.5082 | 0.4914 | 0.0003192 | 0.0005563 | 0.9861 | 0.8484 |
| Chr10 |  | 0.5418 | 0.4552 | 0.003015 | 0.07373 | 0.01390 | 0.1516 |
| CEU10 TH | 2 | 0.5344 | 0.4626 | 0.003046 | 0.000 | 0.6843 | 0.6056 |
| Chr10 |  | 0.5502 | 0.4491 | 0.0006450 | 0.01287 | 0.3157 | 0.3944 |

#### Visualizing model fit

We examined the fit of the data to each model using quantile-quantile (QQ) plots, where the quantile of the observed read counts are plotted against the quantile of the estimated read counts. For the model with two components applied to the reference dataset, the reference and error counts fit the expected values (Figure 1A and supplementary figures).

**Figure 1.**
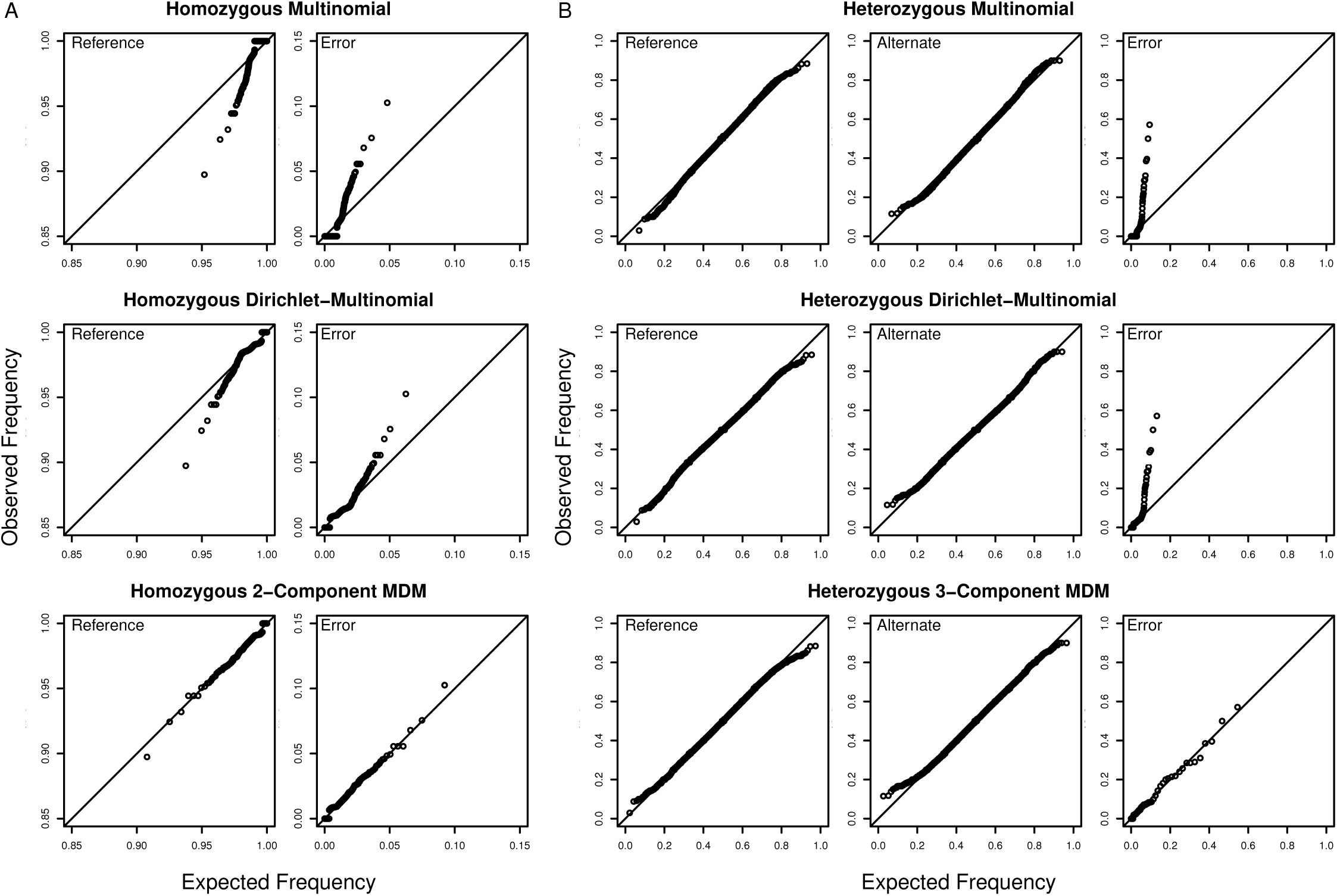
Mixtures of Dirichlet-multinomials provide the best fits to genomic datasets. QQ plots evaluate the fit of three different models to (a) the CHM1 chromosome 21 RD dataset. (b) the CEU 2013 chromosome 21 TH dataset. The quantiles of the observed read count frequencies are calculated from the datasets, and the quantiles of the expected read count frequencies are estimated from the fitted model. A model that fits the data well produces points that fall along the diagonal.

### 3.2 Diploid dataset — Heterozygous

We examined the same two regions in an individual (NA12878 in the CEU trio), along with her parents (NA12891 and NA12892), using data from the 1000 Genomes Project (1000 Genomes Project Consortium *et al.*, 2010, 2015). In order to investigate the impact of sequencing technology on parameter estimates from our model, we repeated our analysis for each of four datasets produced during different stages of the Project. As potential heterozygotes present the greatest challenge to model the distribution we only focused on these sites. Specifically, we identified potential heterozygous sites using the Samtools mpileup function and BCFtools call function on NA12878 alone and using the trio caller Li *et al.* (2009a). Sites were only included in the potential heterozygote (PH) dataset if they were called by both individual and trio methods. This ensured that NA12878 contained enough information to call heterozygote, and it was consistent with the parents. We filtered the PH dataset to include only sites identified as SNPs by the 1000 Genomes Project. We considered these sites to be true heterozygous sites (TH dataset). The number of sites and the proportion of true heterozygous sites are summarized in Table 2.

**Table 2.** Number of heterozygous sites identified by different methods and the number of true heterozygous sites from 1000 Genomes Project.

| Dataset | Individual caller only | Trio caller only | Both callers (PH) | True heterozygotes (TH) |
| --- | --- | --- | --- | --- |
| CEU13 Chr21 | 143 | 1604 | 40956 | 30120 |
| CEU13 Chr10 | 1652 | 2645 | 38542 | 36590 |
| CEU12 Chr21 | 151 | 1171 | 38180 | 29983 |
| CEU12 Chr10 | 106 | 880 | 40144 | 36818 |
| CEU11 Chr21 | 145 | 1190 | 38107 | 29991 |
| CEU11 Chr10 | 114 | 867 | 40156 | 36825 |
| CEU10 Chr21 | 6447 | 4971 | 31773 | 28197 |
| CEU10 Chr10 | 194 | 1173 | 37108 | 35062 |

#### Best fitting models

We fit eight models to each of the 16 CEU datasets (two genomic regions, four release years, two levels of filters): a multinomial, multinomial with reference bias, a DM, and MDMs with two to six components. As expected, the addition of model components increased the likelihood of the MDM model for all cases (Supplementary tables).

For the TH datasets, the best-fitting models had either two or three components depending on the run year. The majority of the sites (88–99%) were assigned to one component in the model (Table 1 and supplementary tables). The major component of the model for each dataset (the component with the highest proportion of sites) had little overdispersion (φ = 0–0.0006).

For the 2011, 2012, and 2013 datasets, the major component had an approximately equal proportion of reference and alternative alleles (49%0–51%), and a relatively small error term (< 0.1%). Thus, the majority of sites fall into a component that is well approximated by a binomial distribution with equal probabilities of reference and alternative alleles. The best-fitting model for the 2010 genome sequence has a slightly larger error term (0.2% and 0.3%), and the reference and alternate terms are 53% and 46% respectively. For the datasets with a two component model, the minor component is similar to the major component but with greater overdispersion (φ = 0.060–0.1).For models with three components, the minor components had an elevated proportion of one of the reference, alternate, or error terms, and greater overdispersion. For instance, CEU2013 chromosome 21 has φ = 0.0656 and π_error_ = 0.278 for the third component.

For the PH dataset, the best fitting model had three to six components (Supplementary tables). The major component of the model contained between 71% and 95% of sites for all years. As with the TH dataset, the major components all had little overdispersion (φ = 0–0.0014). Even when models with greater than four components were favored by BIC, the additional components contained a very small proportion of the data (< 1%), and frequently produced estimates of sequencing error very close to zero. Thus, a model with more than three components was likely overfitting the data.

#### Visualizing model fit

When we examined the QQ plot for the MDM model with the lowest BIC, all three terms (reference allele,alternative allele, and error term) fit closely to the expected values (Figure 1B and supplementary figures). The error term in particular fits the model better than multinomial and multinomial-with-reference-bias models.

#### Copy number variants and low complexity regions

We assigned each site from the eight PH datasets to a component in the best-fitting model based on the site likelihood. Repetitive regions of the genome (e.g. SINEs, ALUs, LINEs, LTRs, and retroposons),low-complexity regions (LCRs), and copy-number variable genes (CNVs) are known to have higher genotype error rate (Li, 2014; Malhis and Jones, 2010). The read counts of these sites are likely to deviate from the expected distributions and differ from other genomic regions. Therefore, we predict that these sites are more likely to be in the minor component.

The minor component of the best-fitting model for each dataset was enriched for CNVs, repetitive regions, and LCRs. In chromosome 21, the proportion of CNVs is 3.7%–6.7% in the minor component and 0.5%–1.5% in the major component. In chromosome 10, the proportion of CNVs is 0.3%–0.5% in the minor component and 0.2%–0.3% in the major component. Fisher’s exact tests show a significantly higher proportion of CNVs sites in the minor component than major component for all datasets (*p* < 0.05) except for CEU10 chromosome 10 (p = 0.198). In the minor component, 53.5–60.7% of sites were in repetitive regions/LCRs, compare to 45.9–51.1% in the major component. Fisher’s exact tests show a significantly higher proportion of repetitive regions/LCRs in the minor component (*p* < 1 × 10^−5^; Table 3).

**Table 3.** The minor components have a higher percentage of copy number variable regions (CNVs) and repetitive/low complexity regions (LCRs). The number and percent (%) of CNV regions, and LCRs are shown for the major and minor components (combined) for the best fit model for each CEU dataset. *p*-value was calculated for the difference between the proportion of CNVs or LCRs in each component. These *p*-values are not corrected for multiple comparison tests.

| Dataset | | Non-CNV/CNV | % | $p$-value | Non-LCR/LCR | % | $p$-value |
| --- | --- | --- | --- | --- | --- | --- | --- |
| CEU13 | Major | 26924 / 415 | 1.5 | 4.11e-143 | 13362 / 13977 | 51.1 | 2.91e-48 |
| Chr21 | Minor | 10851 / 773 | 6.7 |  | 4746 / 6878 | 59.2 |  |
| CEU13 | Major | 32012 / 94 | 0.3 | 0.00419 | 16043 / 16063 | 50.0 | 5.39e-06 |
| Chr10 | Minor | 5864 / 32 | 0.5 |  | 2756 / 3140 | 53.3 |  |
| CEU12 | Major | 26891 / 193 | 0.7 | 4.54e-87 | 13362 / 13722 | 50.7 | 2.67e-57 |
| Chr21 | Minor | 7760 / 331 | 4.1 |  | 3178 / 4913 | 60.7 |  |
| CEU12 | Major | 32560 / 90 | 0.3 | 0.00158 | 16204 / 16446 | 50.4 | 2.08e-16 |
| Chr10 | Minor | 7035 / 37 | 0.5 |  | 3129 / 3943 | 55.8 |  |
| CEU11 | Major | 26858 / 191 | 0.7 | 1.11e-88 | 13344 / 13705 | 50.7 | 3.11e-54 |
| Chr21 | Minor | 7743 / 333 | 4.1 |  | 3194 / 4882 | 60.5 |  |
| CEU11 | Major | 32539 / 89 | 0.3 | 0.00101 | 16195 / 16433 | 50.4 | 3.19e-16 |
| Chr10 | Minor | 7060 / 38 | 0.5 |  | 3144 / 3954 | 55.7 |  |
| CEU10 | Major | 21968 / 109 | 0.5 | 3e-90 | 11936 / 10141 | 45.9 | 3.62e-71 |
| Chr21 | Minor | 8518 / 326 | 3.7 |  | 3790 / 5054 | 57.1 |  |
| CEU10 | Major | 26380 / 49 | 0.2 | 0.198 | 14305 / 12124 | 45.9 | 1.68e-53 |
| Chr10 | Minor | 10197 / 26 | 0.3 |  | 4617 / 5606 | 54.8 |  |

## 4 Discussion

In this paper we have shown that standard models inaccurately represent the distribution of reads for homozygotes and heterozygotes. For every dataset we considered, our MDM approach provided a better fit to NGS data than any single-component model. Indeed, the best-fitting models generated for a haploid human cell line contained two components, demonstrating that NGS datasets generated from relatively simple biological samples (i.e. no true heterozygotes and a high quality reference genome) can benefit from the approach we describe here. Similarly, our MDM model provide a better fit to more complex data, including potentially heterozygous sites in data arising from the 1000 Genomes Project. In order to take into account the excess variations in the PH dataset, the MDM models fit more complex models to the PH dataset (3–6 components) than TH dataset (2–3 components).

### 4.1 Overdispersion and reference bias are not universal in NGS data

Previous work on the statistical properties of NGS data have emphasized the presence of reference bias due to errors in read mapping (Degner *et al.*, 2009; Krawitz *et al.*, 2010;Heinrich *et al.*, 2012), and overdispersion due to correlated errors during library preparation and sequencing (Ramu *et al.*, 2013). Both these processes would move the distribution of reads in an NGS experiment away from a standard binomial distribution with probabilities reflecting the underlying genotype. Our results demonstrate that not all sites in an NGS experiment are subject to these processes. The best-fitting model for every dataset we considered contained a major component with relatively little overdispersion or allelic bias. Sites that fall in these components are thus well approximated by a binomial. On the other hand, a substantial minority of sites in all cases fall into components that do display allelic bias, a high rate of apparent sequencing error, or overdispersion. Our mixture model approach allows us accurately model read distributions for these sites different types of sites, and avoid applying inappropriate model-parameters to the majority of sites.

### 4.2 Copy number variants and low complexity regions

The minor components of our MDM models are enriched for CNVs and low-complexity regions (LCRs). CNVs can cause misalignment of reads in repetitive regions, which is thought to be a cause of genotyping error (Muralidharan *et al.*, 2012). LCRs are particularly prone to misalignment as a result of compositional biases and the alignment of paralogous sequences (Wootton and Federhen, 1996; Frith, 2011; Li, 2014). These genomic regions resulted in misalignment of reads and altered the profile of the read counts for some sites. By automatically partitioning these regions in a separate component of the MDM model we avoid applying inappropriate model-parameters to the majority of sites.

### 4.3 Differences among datasets

Although all of the diploid datasets contained reads produced from Illumina sequencing, a variety of different library preparations and sequencing platforms were used to produce the data. The 2010 data was generated using short paired reads (35bp–67bp) and relatively error-prone sequencing. The 2011 and 2012 datasets contain reads of intermediate length (101bp) mapped to a reference genome designed to lower the rate of misalignment. The 2013 dataset contains the longest reads (250bp) and was produced using a PCR-free library preparation. Our approach was able to fit each of these datasets well, regardless of the library preparations and sequencing technologies.

The parameter values estimated by EM in each dataset reflect the technology used. Not surprisingly, the 2010 dataset appears to have a quite different error profile to other years: the major component of this model has higher overdispersion and stronger reference bias than that of any other dataset. The major components of datasets from 2011 and 2012 fit more closely to the expectation of a binomial distribution with equal frequencies of reference and non-reference alleles. Thus we conclude that advances in sequencing technologies have led to the majority of sites in NGS datasets better fitting basic expectations of how alleles are amplified, sequenced, and mapped.

However, while we expected the 2013 dataset to be the “cleanest” dataset due to reducing polymerase base misincorporation errors, this was not the case. Nevertheless, the 95% confidence interval of the overdispersion parameter, shows no significant difference between most of the datasets. The only exception is chromosome 21 from CEU13, where the value of the overdispersion parameter is significantly higher (Figure 2). Thus, we suggest reconsidering the effectiveness of the PCR-free approach.

**Figure 2.**
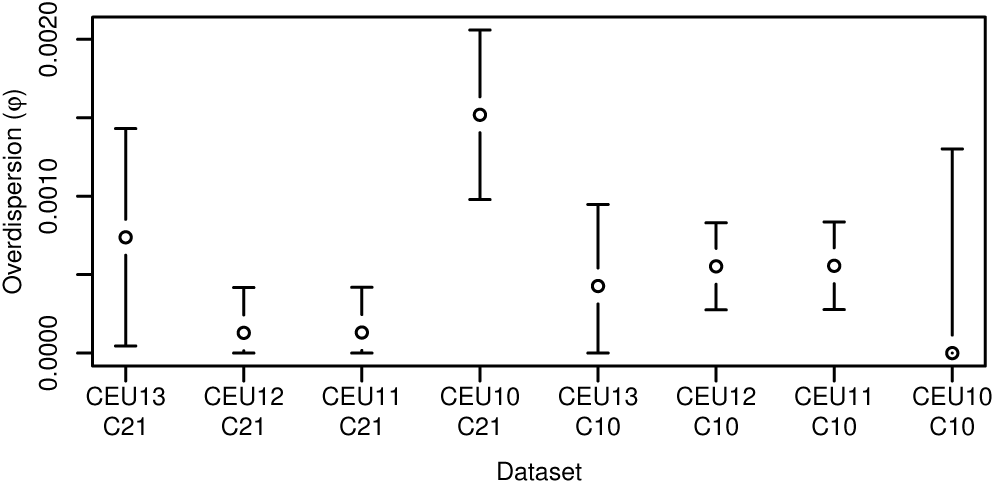
Confidence interval for φ from the major component in the two components model for different TH datasets. Most of the estimated φ overlap in chromosome 21 and all of them in chromosome 10.

We conclude that read distributions for a majority of sites fit a basic binomial distribution. This contrasts with recent approaches to improve genotyping that fit a more complex beta-binomial. Only a minority of sites are subject to processes that drive read count distributions away from a binomial distribution. Correctly modeling these sites separately should increase genotyping accuracy. The approach we describe here will be included in future versions of the mutation-detecting software DeNovoGear and accuMUlate (Ramu *et al.*, 2013; Long *et al.*, 2016).

## Acknowledgements

The authors declare that they have no competing interests.

## References

1000 Genomes Project Consortium, Abecasis, G. R., Altshuler, D.,Auton, A., Brooks, L. D., Durbin, R. M., Gibbs, R. A., Hurles, M. E., and McVean, G. A.(2010). A map of human genome variation from population-scale sequencing. Nature, 467(7319), 1061–1073.

1000 Genomes Project Consortium, Auton, A., Brooks, L. D., Durbin, R. M., Garrison, E. P., Kang, H. M., Korbel, J. O., Marchini, J. L., McCarthy, S., McVean, G. A., and Abecasis, G. R. (2015). A global reference for human genetic variation. Nature, 526(7571), 68–74.

Awadalla, P., Gauthier, J., Myers, R. A., Casals, F., Hamdan, F. F., Griffing, A. R., Côté, M., Henrion, E., Spiegelman, D., Tarabeux, J., Piton, A., Yang, Y., Boyko, A., Bustamante, C., Xiong, L., Rapoport, J. L., Addington, A. M., DeLisi, J. L. E., Krebs, M.-O., Joober, R., Millet, B., Fombonne, E., Mottron, L., Zilversmit, M., Keebler, J., Daoud, H., Marineau, C., Roy-Gagnon, M.-H., Dubé, M.-P., Eyre-Walker, A., Drapeau, P., Stone, E. A., Lafrenière, R. G., and Rouleau, G. A. (2010). Direct measure of the de novo mutation rate in autism and schizophrenia cohorts. Am J Hum Genet, 87(3), 316–324.

Cartwright, R. A., Hussin, J., Keebler, J. E. M., Stone, E. A., and Awadalla, P. (2012). A family-based probabilistic method for capturing de novo mutations from high-throughput short-read sequencing data. Stat Appl Genet Mol Biol, 11(2).

Degner,J.F., Marioni, J. C., Pai, A.A., Pickrell, J. K., Nkadori, E., Gilad, Y., and Pritchard, J.K. (2009). Effectofread-mapping biasesondetecting allele-specific expression from RNA-sequencing data. Bioinformatics, 25(24), 3207–3212.

DePristo, M. A., Banks, E., Poplin, R., Garimella, K. V., Maguire, J. R., Hartl, C., Philippakis, A. A., del Angel, G., Rivas, M. A., Hanna, M., and et al. (2011). A framework for variation discovery and genotyping using next-generation DNA sequencing data. Nat Genet, 43(5), 491–498.

Farrer, R. A., Henk, D. A., MacLean, D., Studholme, D. J., and Fisher, M. C. (2013). Using false discovery rates to benchmark SNP-callers in next-generation sequencing projects. Sci Rep, 3.

Fox, E. J., Reid-Bayliss, K. S., Emond, M. J., and Loeb, L. A. (2014). Accuracy of next generation sequencing platforms. Next Gener Seq Appl, 1.

Frith, M. C. (2011). Gentle masking of low-complexity sequences improves homology search. PloS One, 6(12), e28819.

Goldstein, D. B., Allen, A., Keebler, J., Margulies, E. H., Petrou, S., Petrovski, S., and Sunyaev, S. (2013). Sequencing studies in human genetics: design and interpretation. Nat Rev Genet, 14(7), 460–470.

Goya, R., Sun, M. G., Morin, R. D., Leung, G., Ha, G., Wiegand, K. C., Senz, J., Crisan, A., Marra, M. A., Hirst, M., et al. (2010). SNVMix: predicting single nucleotide variants from next-generation sequencing of tumors. Bioinformatics, 26(6), 730–736.

Harismendy, O., Ng, P. C., Strausberg, R. L., Wang, X., Stockwell, T. B., Beeson, K. Y., Schork, N. J., Murray, S. S., Topol, E. J., Levy, S., and Frazer, K. A. (2009). Evaluation of next generation sequencing platforms for population targeted sequencing studies. Genome Biol, 10(3), R32.

Heinrich, V., Stange, J., Dickhaus, T., Imkeller, P., Krüger, U., Bauer, S., Mundlos, S., Robinson, P. N., Hecht, J., and Krawitz, P. M. (2012). The allele distribution in next-generation sequencing data sets is accurately described as the result of a stochastic branching process. Nucleic Acids Res, 40(6), 2426–2431.

Josephidou, M., Lynch, A. G., and Tavaré, S. (2015). multiSNV: a probabilistic approach for improving detection of somatic point mutations from multiple related tumour samples. Nucleic Acids Res, 43(9), e61.

Karolchik, D., Hinrichs, A. S., Furey, T. S., Roskin, K. M., Sugnet, C. W., Haussler, D., and Kent, W. J. (2004). The UCSC Table Browser data retrieval tool. Nucleic Acids Res, 32(Database issue), D493–D496.

Koboldt, D. C., Steinberg, K. M., Larson, D. E., Wilson, R. K., and Mardis, E. R. (2013). The next-generation sequencing revolution and its impact on genomics. Cell, 155(1), 27–38.

Krawitz, P., Rödelsperger, C., Jäger, M., Jostins, L., Bauer, S., and Robinson, P. N. (2010). Microindel detection in short-read sequence data. Bioinformatics, 26(6), 722–729.

Li, H. (2011). A statistical framework for SNP calling, mutation discovery, association mapping and population genetical parameter estimation from sequencing data. Bioinformatics, 27(21), 2987–2993.

Li, H. (2014). Toward better understanding of artifacts in variant calling from high-coverage samples. Bioinformatics, 30(20), 2843–2851.

Li, H., Ruan, J., and Durbin, R. (2008a). Mapping short DNA sequencing reads and calling variants using mapping quality scores. Genome Res, 18(11), 1851–1858.

Li, H., Handsaker, B., Wysoker, A., Fennell, T., Ruan, J., Homer, N., Marth, G., Abecasis, G., Durbin, R., and 1000 Genome Project Data Processing Subgroup (2009a). The sequence alignment/map format and SAMtools. Bioinformatics, 25(16), 2078–2079.

Li, R., Li, Y., Kristiansen, K., and Wang, J. (2008b). SOAP: short oligonucleotide alignment program. Bioinformatics, 24(5), 713–714.

Li, R., Li, Y., Fang, X., Yang, H., Wang, J., Kristiansen, K., and Wang, J. (2009b). SNP detection for massively parallel whole-genome resequencing. Genome Res, 19(6), 1124–1132.

Long, H., Winter, D. J., Chang, A. Y.-C., Sun, W., Wu, S. H., Balboa, M., Azevedo, R. B., Cartwright, R. A., Lynch, M., and Zufall, R. A. (2016). Low base-substitution mutation rate in the ciliate Tetrahymena thermophila. Genome Biol Evol.

Malhis, N. and Jones, S. J. M. (2010). High quality SNP calling using Illumina data at shallow coverage. Bioinformatics, 26(8), 1029–1035.

Mills, R. E., Walter, K., Stewart, C., Handsaker, R. E., Chen, K., Alkan, C., Abyzov, A., Yoon, S. C., Ye, K., Cheetham, R. K., Chinwalla, A., Conrad, D. F., Fu, Y., Grubert, F., Hajirasouliha, I., Hormozdiari, F., Iakoucheva, L. M., Iqbal, Z., Kang, S., Kidd, J. M., Konkel, M. K., Korn, J., Khurana, E., Kural, D., Lam, H. Y. K., Leng, J., Li, R., Li, Y., Lin, C.-Y., Luo, R., Mu, X. J., Nemesh, J., Peckham, H. E., Rausch, T., Scally, A., Shi, X., Stromberg, M. P., Stütz, A. M., Urban, A. E., Walker, J. A., Wu, J., Zhang, Y., Zhang, Z. D., Batzer, M. A., Ding, L., Marth, G. T., McVean, G., Sebat, J., Snyder, M., Wang, J., Ye, K., Eichler, E. E., Gerstein, M. B., Hurles, M. E., Lee, C., McCarroll, S. A., Korbel, J. O., and, G. P. (2011). Mapping copy number variation by population-scale genome sequencing. Nature, 470(7332), 59–65.

Muralidharan, O., Natsoulis, G., Bell, J., Newburger, D., Xu, H., Kela, I., Ji, H., and Zhang, N. (2012). A cross-sample statistical model for SNP detection in short-read sequencing data. Nucleic Acids Res, 40(1), e5.

Peng, G., Fan, Y., Palculict, T. B., Shen, P., Ruteshouser, E. C., Chi, A.-K., Davis, R. W., Huff, V., Scharfe, C., and Wang, W. (2013). Rare variant detection using family-based sequencing analysis. Proc Natl Acad Sci U S A, 110(10), 3985–3990.

Ramu, A., Noordam, M. J., Schwartz, R. S., Wuster, A., Hurles, M. E., Cartwright, R. A., and Conrad, D. F. (2013). DeNovoGear: de novo indel and point mutation discovery and phasing. Nat Methods, 10(10), 985–987.

Sayed, S., Langdon, D. R., Odili, S., Chen, P., Buettger, C., Schiffman, A. B., Suchi, M., Taub, R., Grimsby, J., Matschinsky, F. M., and Stanley, C. A. (2009). Extremes of clinical and enzymatic phenotypes in children with hyperinsulinism caused by glucokinase activating mutations. Diabetes, 58(6), 1419–1427.

Tvedebrink, T. (2010). Overdispersion in allelic counts and u-correction in forensic genetics. Theor Popul Biol, 78(3), 200–210.

Wall, J. D., Tang, L. F., Zerbe, B., Kvale, M. N., Kwok, P.-Y., Schaefer, C., and Risch, N. (2014). Estimating genotype error rates from high-coverage next-generation sequence data. Genome Res, 24(11), 1734–1739.

Warr, A., Robert, C., Hume, D., Archibald, A. L., Deeb, N., and Watson, M. (2015). Identification of low-confidence regions in the pig reference genome (Sscrofa10.2). Front Genet, 6, 338.

Wootton, J. C. and Federhen, S. (1996). Analysis of compositionally biased regions in sequence databases. Methods Enzymol, 266.

